# Conservation agriculture reduces climate change impact of a popcorn and wheat crop rotation

**DOI:** 10.1101/2022.12.22.521544

**Authors:** Maria Vittoria Guidoboni, Annie Duparque, Joachim Boissy, Jean-Christophe Mouny, Julie Auberger, Hayo M.G. van der Werf

## Abstract

Urgent action is needed to ensure humanity’s future under climate change. Agriculture faces major challenges as it is both influenced by and contributes to climate change. Conservation agriculture reduces greenhouse gas (GHG) emissions and sequesters carbon (C) in the soil due to practices such as reduced tillage and planting of cover crops. This study assessed effects of an innovative conservation agriculture popcorn (*Zea mays*) and wheat (*Triticum aestivum*) crop rotation in south-western France on soil C sequestration, GHG emissions and several environmental impacts. Two complementary approaches were used: i) a comparison based on field data and expert judgement to assess short-term effects and ii) modelling of three scenarios to quantify long-term outcomes. In both approaches Life cycle assessment (LCA) was used to compare popcorn and wheat rotations. The conventional rotation used ploughing, and its soil was bare between wheat harvest and popcorn sowing. Conservation agriculture used reduced tillage, cover crops, and compost of green waste. Impacts of compost production were allocated mainly to its waste treatment function, based on waste treatment cost and compost price. Simulation modelling of soil C was used to estimate the amount of C sequestered by the conservation and conventional crop rotations. LCA was combined with soil C modelling over 100 years to assess the long-term climate change impact of three scenarios for the popcorn and wheat rotation. Mean annual C sequestration and net climate change impact were -0.24 t/ha and 3867 kg CO_2_-eq./ha, respectively, for the conventional rotation and 0.91 t/ha and 434 kg CO_2_-eq./ha, respectively, for the conservation rotation. The climate change impact of the conservation rotation depended strongly on the allocation of composting impacts between the waste treatment and compost production functions. Compared to the conventional rotation, the conservation rotation had a lower marine eutrophication impact (−7%) but higher impacts for terrestrial acidification (+9%), land competition (+3%), and cumulative energy demand (+2%). Modelling over 100 years revealed that at near soil C equilibrium, a conventional scenario lost 9% of soil C, whereas conservation agriculture scenarios gained 14% (only cover crop) and 26% of soil C (cover crop + compost). Conservation agriculture resulted in soil C sequestration over several decades, until a new soil C equilibrium was reached.

**Highlights:**

- Conservation and conventional popcorn and wheat crop rotations were compared
- Coupling of LCA and soil carbon modelling allowed for comprehensive assessment
- Conservation agriculture sequestered carbon in the soil
- Conservation agriculture strongly reduced climate change impact
- Compost impact-allocation choices strongly influenced potential impacts

## 1. Introduction

The Intergovernmental Panel on Climate Change (IPCC, 2021) has reiterated the need for urgent action to ensure the future of humanity under climate change. To remain within the target of no more than 1.5°C of warming, urgent action at unprecedented scales and across all sectors is required within the next 10 years. According to IPCC (2014), 24% of total greenhouse gas (GHG) emissions worldwide are caused by the agricultural sector. Consequently, agriculture has a major role in meeting the 1.5°C target through lowering its GHG emissions and sequestering carbon dioxide (CO_2_) in the soil. At the same time, agricultural systems need to adapt to climate change, and produce sufficient food for a growing world population, while preserving biodiversity and water and soil resources.

Soil carbon (C) sequestration, which represents up to 90% of the mitigation potential for the global agricultural sector, can contribute to reducing net anthropogenic GHG emissions, while preserving and restoring soil health by maintaining and increasing soil organic matter, thus helping to increase production resilience and food security (Smith et al., 2014). Conservation agriculture includes a variety of agricultural practices, such as reduced tillage, permanent organic soil cover with crop residues and/or cover crops, and crop rotation (Hobbs et al., 2008; Rusinamhodzi et al., 2011).

Conservation agriculture addresses several environmental and management issues. Planting of cover crops can improve soil quality by providing a source of substrate for microbial activity, which enhances soil C and nitrogen (N) cycles (Mbuthia et al., 2015). Soil erosion decreases when soils are less disturbed by tillage and are enriched with biomass from cover crops and organic fertilisation (Lal, 2015). Cover crops also increase N-use efficiency: they take up soil N left by the previous crop, thus decreasing N leaching and providing N from cover crop residues to the next crop (Hartwig and Ammon, 2002). Finally, conservation agriculture usually has lower production costs than conventional agriculture, due to reduced use of inputs (e.g. fuel) and less time spent in the field (Giller et al., 2015; Hobbs, 2008).

Conservation agriculture is not common in popcorn (*Zea mays*) and wheat (*Triticum aestivum*) rotations in France. In current conventional popcorn and wheat rotations, the soil is ploughed in autumn, remains bare in winter, and only mineral fertiliser is applied. This type of management is prone to soil erosion and nitrate (NO_3_) leaching. It also results in low stocks of soil organic matter, which negatively influences soil structure and soil water retention. The Nataïs company, Europe’s main popcorn producer, started transitioning towards conservation agriculture approximately two decades ago (Bodoville, 2014), with reduced tillage, inclusion of cover crops, and application of organic fertiliser.

Several studies have highlighted the importance of soil organic C (SOC) dynamics when estimating the C footprint of agricultural systems, particularly when conservation agriculture practices are used (Plaza-Bonilla et al., 2018; Yao et al., 2017). These studies have shown that planting cover crops and using reduced or no tillage are good strategies to increase soil C content and reduce net GHG emissions of cropping systems. Although these practices tend to increase in-field N_2_O emissions, due to the incorporation of cover crop residues, and may cause additional CO_2_ emissions related to fuel combustion, conservation agriculture has lower net GHG emissions than conventional agriculture (Prechsl et al., 2017; Yao et al., 2017). Similarly, Prechsl et al. (2017) found that using organic fertiliser (cattle slurry) rather than mineral N fertiliser in a 6-year rotation with cover crops in Switzerland emitted more in-field N_2_O. However, the GHG emissions associated with production of mineral N fertiliser tipped the net GHG emissions in favour of organic fertilisation.

The objectives of this study were to (i) assess short-term effects of an innovative conservation agriculture popcorn and wheat crop rotation on SOC sequestration, GHG emissions, and several environmental impacts, and (ii) to quantify the potential mid- to long-term benefits of conservation agriculture for SOC sequestration and environmental impacts.

## 2. Materials and methods

### 2.1 Life cycle assessment

We used life cycle assessment (LCA) to assess environmental impacts. LCA is a methodological framework that estimates potential environmental impacts of a product by quantifying the resources consumed and pollutant emissions to the environment in the course of its life cycle (ISO, 2006a, 2006b). Conservation agriculture differs from conventional agriculture both with respect to input use (e.g. tractors, seed for cover crops) and with respect to its sequestration of SOC. LCA allows a whole system comparison of conventional and conservation agriculture by considering simultaneously off-farm resource use and emissions of GHG and other pollutants (associated with inputs), on-farm resource use and emissions of GHG and other pollutants (e.g. from diesel combustion) and SOC changes corresponding to CO_2_ emissions or sequestration.

#### 2.1.1 Goal and scope definition

The goal of the study was to compare potential environmental impacts of conventional and conservation agriculture in a popcorn and wheat crop rotation. Two complementary approaches were used: i) comparing the two rotations based on field data (for conservation agriculture) and expert judgement (for conventional agriculture) to assess short-term effects and ii) modelling three agricultural scenarios to quantify long-term outcomes.

##### Comparison of two crop rotations to assess short-term effects

Field data (i.e. farmer practices and yields) on the conservation agriculture crop rotation was collected on the Nataïs commercial farm (43°52’ North, 0°89’ East) in the Gers administrative department (south-western France) during two rotations: 2017-2019 and 2018-2020. Results for each crop were calculated by averaging results for five wheat or popcorn fields. With slopes of 4-12%, the Nataïs farm’s soils are prone to erosion, even though they contain 25-30% clay. The study site has a temperate oceanic climate according to the Köppen-Geiger climate classification (Peel et al., 2007), with mean monthly precipitation of 54 mm and mean annual air temperature of 15.1°C from 2018-2020. The 2018-2020 period was relatively warm, and the 30-year mean air temperature was 13.5°C. Soil texture was mainly clay-limestone. The initial soil organic matter content (0-30 cm) at the experimental sites was 1.8-2.1%, and the initial SOC stock was 10.3-12.0 g/kg.

In the conservation agriculture rotation, popcorn received compost of green waste along with mineral fertiliser, while winter wheat received only mineral fertiliser. Green waste consisted mainly of grass, leaves, and branch cuttings. Popcorn was sown according to a technique called “green-tillage” as described hereafter. For the 2017-2019 rotation, winter wheat was sown in October 2017. In July 2018, after the wheat harvest, stubble was tilled twice, at 5 and 10 cm. In mid-September 2018, the soil was tilled with a spring-toothed harrow, to a depth of 20 cm on the future popcorn row with one set of teeth, and to a depth of 8 cm with a second set of teeth. Some of the soil of the future inter row was moved to the future row to form a ridge. In the same operation, winter fava bean (98.6%) and phacelia (1.4%) was sown to establish a cover crop in the future inter-row. During winter, the soil of the ridge settled, which created a seedbed for the popcorn. In April 2019, herbicide application killed the cover crop. Subsequently, popcorn was sown on the ridges using a machine that flattened the cover crop and simultaneously applied fertiliser, insecticide, and molluscicide on the row. Sowing popcorn on ridges favoured germination due to higher soil temperature. After harvesting the popcorn in October 2019, winter wheat was directly sown and mineral fertiliser was applied. For the 2018-2020 rotation, the operations were identical, but two cover crops were grown: a sorghum cover crop was sown at the beginning of July, and chopped and left on the field in mid-September before sowing the fava bean-phacelia cover crop.

Field data for the conventional agriculture rotation reflected dominant farming practices for popcorn and winter wheat rotations in the Gers department according to expert judgment (S. Hypolite, Agro d’Oc, pers. comm.) and are presented hereafter. Popcorn and wheat yields were assumed to be the same as those in the conservation agriculture rotation, and both crops received only mineral fertilisers. For popcorn, the soil was ploughed in autumn and superficially tilled twice in spring. After the popcorn harvest, stubble was tilled at a depth of 10 cm before sowing the winter wheat. No cover crops were grown. All other farming practices were the same as in the conservation agriculture system.

##### Comparison of three scenarios for long-term outcomes

In the scenario-modelling approach, a conventional scenario (Conv) and two conservation scenarios (Cons_CC and Cons_full) were compared. (Table 1). The Conv scenario corresponded to the conventional popcorn and wheat rotation described previously. The two conservation scenarios were based on Nataïs’ conservation agriculture rotation. The Cons_CC scenario used sorghum and fava bean + phacelia as cover crops (as in the 2018-2020 cropping system), whereas the Cons_full scenario used these cover crops as well as 6.5 t/ha of compost of green waste (as in the 2017-2019 cropping system). For all scenarios, yields were set at 6.8 t/ha for winter wheat and 6.3 t/ha for popcorn, based on expert judgement for long-term average yield level of these crops.

**Table 1.** Scenarios of conventional and conservation popcorn and wheat crop rotations

| Scenario | Agricultural system | Cover crops | Compost applied (t/ha) |
| --- | --- | --- | --- |
| Conv | Conventional | Absent | 0 |
| Cons_CC | Conservation | sorghum and fava bean + phacelia | 0 |
| Cons_full | Conservation | sorghum and fava bean + phacelia | 6.5 |

The system boundaries of the crop rotations for both short- and long-term approaches began with the extraction of resources and ended at the farm gate. Processes included the production of all inputs and all processes related to crop production: (1) soil cultivation, sowing, weed control, fertilisation, pest and pathogen control, harvest; (2) machines and the buildings or areas used to park them; (3) seeds for cash and cover crops, fertilisers including compost, pesticides, water for irrigation and fuel, as well as their transport to the farm; (4) direct emissions of fuel combustion, tire abrasion and pollutant emissions in the field. Post-harvest processes, such as drying, sorting and storage were not included. The function considered was “land management”, defined as the occupation of agricultural and non-agricultural land for a given amount of time. The land occupied included both “direct” land (on-farm) and “indirect” land (off-farm, e.g. to produce seeds for sowing). The functional unit was 1 ha of land occupied for 1 year (i.e. ha.year).

#### 2.1.2 Life cycle inventory

Life cycle inventories (LCIs) were calculated using the MEANS-InOut web application v3.0, which is a customized agricultural LCA tool that generates LCIs of agricultural production systems (Auberger et al., 2018). It contains forms to guide data entry and includes a reference data set for the pollutant emissions and resource use of main inputs of agri-food systems, analytical models to estimate direct pollutant emissions and resource use, and an export function that generates LCI files ready to be imported into LCA software to calculate impact indicators. Databases used for background processes (e.g. production of fertilizer, irrigation infrastructure), were AGRIBALYSE v3.0.1 and ecoinvent v3.5.

Emissions to the air (ammonia (NH_3_), nitrous oxide (N_2_O), N oxides (NO_x_), CO_2_), water (NO_3_, phosphorus (P), phosphate (PO_4_), cadmium (Cd), chrome (Cr), copper (Cu), mercury (Hg), nickel (Ni), lead (Pb), zinc (Zn)) and soil (Cd, Cr, Cu, Hg, Ni, Pb, Zn, pesticides) were calculated using models recommended by the AGRIBALYSE methodology (Koch and Salou, 2020). NH_3_ emissions from organic fertiliser application were modelled using EMEP/EEA 2016 Tier 2 (EMEP/EEA, 2016). N_2_O emissions were modelled using IPCC 2019 Tier 1 (IPCC, 2019). NO_x_ emissions were modelled using EMEP/EEA 2009 Tier 1 (EMEP/EEA, 2009). NO_3_ emissions were modelled according to Tailleur et al. (2012). CO_2_ emissions from fuel combustion and emissions of active substances of pesticides were modelled according to ecoinvent® v2 (Nemecek and Kägi, 2007).

The LCI of the compost of green waste available in the AGRIBALYSE database (Avadi, 2020) attributes all polluting emissions of the composting process to the compost, and thus to the crop on which the compost is applied. From a methodological point of view, this is questionable, as the composting process has two functions: waste treatment and compost production. According to the LCI for compost in the AGRIBALYSE database, producing 1 t of compost requires 3 t of green waste. The cost of treating green waste by composting was assumed to be 60 €/t (ADEME, 2002) (i.e. 180 € per t of compost), and the price of the green waste compost delivered to the Nataïs farm was 14 €/t. Given a total cost of 1 t of compost of 194 €/t, 7.2% of it (i.e. 14/194×100) was for the compost and the rest for the waste treatment process. Consequently, following an approach proposed by Christensen et al. (2017), we used economic allocation to allocate pollutant emissions and resource use between the two functions: 7.2% to compost production and 92.8% to waste treatment.

#### 2.1.3 Impact assessment: LCA impact categories

Impacts were assessed using SimaPro software v9.0.0.35. We calculated climate change, terrestrial acidification, and marine eutrophication impacts according to the ReCiPe 2016 Midpoint (H) characterisation method (Huijbregts et al., 2020). We also calculated land competition impact according to the CML-IA non-baseline method (Guinée, 2002) and cumulative energy demand (Frischknecht et al., 2007).

### 2.2 Estimating changes in soil organic carbon stock

Most LCA studies of agricultural production systems assume that farming practices remain constant over time, and thus that the SOC stock remains constant. This assumption is often reasonable, but not when considering a change in farming practices that increases the input of organic C to the soil, such as the introduction of cover crops and organic fertiliser. When such practices are implemented, the increase in SOC stock needs to be included in the LCI, as it represents an absorption of CO_2_ from the atmosphere (i.e. a negative emission).

The change in SOC stock for the comparison of two rotations was estimated using the AMG model (Clivot et al., 2019), which considers three compartments: fresh exogenous organic C, active SOC, and stable SOC. The sources of fresh organic C are organic fertilisers and crop residues. The active SOC compartment is fed by fresh organic C inputs and influenced by annual mineralisation, while stable SOC is considered to be completely inert over the short and mid-terms (i.e. its turnover time is millenary). The AMG model represents the generally accepted fact that the SOC pool is heterogeneous. The AMG model has three main parameters: the humification coefficient, which is the conversion factor from fresh organic C inputs into humified SOC; the annual rate of SOC mineralisation; and the initial proportion of stable C in the initial SOC stock. The humification coefficient depends only on the nature of fresh organic C inputs, whereas the annual rate of SOC mineralisation depends on the soil (i.e. clay and lime contents, pH, C:N ratio of soil organic matter), soil tillage (type and depth), and meteorological conditions (i.e. mean annual air temperature and water balance). The model runs on a yearly basis at the field level to estimate SOC stock at a depth of 0-30 cm.

To estimate the amount of crop biomass left on the field after harvest, crop yield is converted into above-ground biomass using a crop-specific harvest index. Below-ground biomass is estimated using an allometric relationship with aerial biomass (Baret et al., 1992). Total (above- and below-ground) biomass minus crop yield equals the amount of biomass that provides fresh organic C inputs. In the comparison of the conservation and the conventional agriculture crop rotation field data for crop yields were available for conservation agriculture but not for conventional agriculture. We assumed that crop yields of the latter were identical to those of the former. This is not necessarily true, we therefore conducted a sensitivity analysis assessing the effects of a 10% increase or decrease of the yield of conventional crops on carbon storage and subsequent environmental impacts.

For the comparison of three scenarios, changes in the SOC stock over 100 years were estimated using the SIMEOS-AMG tool (Duparque et al., 2011; 2013), which is a decision-support tool based on the AMG model designed to facilitate simulation of long-term changes in SOC. Model description and model inputs and outputs for AMG and SIMEOS-AMG are presented in the supplementary material (Tables S1 and S2).

## 3. Results

### 3.1 Characteristics of crop rotations

Wheat yielded 5.9 and 9.5 t/ha in 2018 and 2019 respectively, whereas popcorn yielded 6.5 and 6.2 t/ha in 2019 and 2020, respectively (Table 2). The fava bean/phacelia cover crop produced 3.1 and 3.9 t dry matter/ha in 2019 and 2020, respectively, whereas the sorghum cover crop produced 3 t dry matter/ha in 2020.

**Table 2.** Yields, inputs, and main direct emissions for conventional and conservation wheat and popcorn crop rotations in 2017-2019 and 2018-2020. WW=Winter Wheat; PC= Popcorn. The year indicates the harvest year. Unless indicated otherwise, inputs and direct emissions of cover crops were attributed to conservation popcorn.

|  |  | Conventional |  |  |  | Conservation |  |  |  |
| --- | --- | --- | --- | --- | --- | --- | --- | --- | --- |
|  | Unit | WW_2018 | PC_2019 | WW_2019 | PC_2020 | WW_2018 | PC_2019 | WW_2019 | PC_2020 |
| Yield, cash crop | t/ha | 5.9 | 6.5 | 9.5 | 6.2 | 5.9 | 6.5 | 9.5 | 6.2 |
| Input/Process |  |  |  |  |  |  |  |  |  |
| Seeds, cash crop | kg/ha | 127 | 15 | 122 | 16 | 127 | 15 | 122 | 16 |
| Seeds, cover crop <sup>1</sup> | kg/ha | - | - | - | - | - | 120 | - | 154 |
| Fava bean/phacelia cover crop dry matter | t/ha | - | - | - | - | - | 3.1 | - | 3.9 |
| Sorghum cover crop dry matter | t/ha | - | - | - | - | - | - | - | 3 |
| Compost of green waste | t/ha | - | - | - | - | - | 6.5 | - | 9.9 |
| N compost of green waste | kg/ha | - | - | - | - | - | 52 | - | 79 |
| N mineral fertiliser | kg/ha | 179 | 190 | 202 | 190 | 179 | 188 | 202 | 182 |
| N total | kg/ha | 179 | 190 | 202 | 190 | 179 | 240 | 202 | 261 |
| P<sub>2</sub>O<sub>5</sub> total | kg/ha | 24 | 50 | 24 | 50 | 24 | 53 | 24 | 66 |
| K<sub>2</sub>O total | kg/ha | 0 | 50 | 0 | 50 | 0 | 89 | 0 | 122 |
| Diesel | kg/ha | 40 | 73 | 40 | 75 | 32 | 68 | 32 | 78 |
| Irrigation water | m<sup>3</sup>/ha | - | 1 659 | - | 2 396 | - | 1 659 | - | 2 396 |
| Pesticide active ingredient | kg/ha | 5.2 | 2.8 | 6.6 | 2.8 | 5.2 | 3.7 | 6.6 | 3.7 |
| Direct emissions |  |  |  |  |  |  |  |  |  |
| To air |  |  |  |  |  |  |  |  |  |
| NH<sub>3</sub> | kg/ha | 18.0 | 24.5 | 19.1 | 22.3 | 18.6 | 27.3 | 20.2 | 26.1 |
| NO | kg/ha | 4.4 | 4.7 | 4.9 | 4.5 | 4.6 | 5.4 | 5.1 | 5.4 |
| N<sub>2</sub>O | kg/ha | 5.7 | 6.3 | 6.6 | 6.2 | 5.7 | 8.3 | 6.6 | 9.7 |
| CO<sub>2</sub> | kg/ha | 129 | 209 | 143 | 414 | 129 | 208 | 143 | 410 |
| To water |  |  |  |  |  |  |  |  |  |
| NO<sub>3</sub> | kg/ha | 133 | 111 | 133 | 111 | 111 | 111 | 111 | 111 |

<sup>1</sup> in 2019 only fava bean and phacelia seeds, in 2020 both fava bean, phacelia and sorghum seeds

Conservation popcorn received 6.5 and 9.9 t/ha of compost in 2019 and 2020, respectively, corresponding to 22% and 30% of total N applied, respectively. Compared to conventional popcorn, conservation popcorn (including the cover crop) used 26% (2019) and 37% (2020) more fertiliser N, 6% less (2019) and 5% more (2020) diesel, and 33% (2019) and 32% (2020) more pesticide active ingredient. Both conservation wheat crops (2018 and 2019) used 20% less diesel than the conventional ones. Yields and amounts of cash-crop seed, irrigation water, mineral fertiliser, and pesticide applied were identical for conventional and conservation wheat crops.

Emissions of NH_3_, NO, and N_2_O were higher for the conservation popcorn crops (including cover crop) than for the conventional ones by 11%, 15%, and 32%, respectively, in 2019 and by 17%, 22%, and 58%, respectively, in 2020, whereas CO_2_ emissions were similar. NH_3_ and NO emissions were higher for conservation wheat crops than for conventional wheat crops by 3% and 4%, respectively, in 2019 and by 6% for both gases in 2019. NO_3_ emissions were 17% lower for conservation wheat crops than for conventional wheat crops.

According to the AMG model, conventional wheat and popcorn sequestered -0.24 t C/ha for both the 2017-2019 and the 2018-2020 crop rotation (Table 3). A 10% decrease or increase of assumed yields of the crops in the conventional rotation resulted in a moderate (−0.33/-0.16 and -0.34/-0.15) variation of these values (Table 3). C sequestration of conservation wheat was similar to that of conventional wheat, while conservation popcorn, due to the cover crop and compost application, sequestered 1.6 (2019) and 2.7 (2020) t C/ha. Mean annual C sequestration was 0.6 and 1.2 t C/ha for the 2017-2019 and 2018-2020 conservation crop rotations, respectively. The difference in annual C sequestration between the two systems was thus 0.83 and 1.47 t C/ha, with C input from compost of 0.46 and 0.69 t/ha, for 2017-2019 and 2018-2020, respectively. For conservation popcorn, percentages of C input from crop residues, cover crop, and compost were 36%, 29%, and 35%, respectively, in 2019 and 23%, 43%, and 34%, respectively, in 2020.

**Table 3.** Carbon (C) balance per crop and its components for wheat and popcorn estimated by the AMG model: organic C input from cash-crop residues (Ccrop), mineralisation of soil C (Cmin), organic C input from cover crops (Ccover), organic C input from compost (Ccomp), and humified C sequestered Cseq) for the conventional and conservation rotations. Values are expressed as t C/ha/year at a depth of 0-30 cm. Values in brackets for Cseq of the conventional crops correspond to the effects of a 10% increase or decrease of their yields.

| Year | Crop | Conventional |  |  | Conservation |  |  |  |  |
| --- | --- | --- | --- | --- | --- | --- | --- | --- | --- |
|  |  | Ccrop | Cmin | Cseq | Ccrop | Ccover | Ccomp | Cmin | Cseq |
| 2017-2018 | Wheat | 0.78 | -1.15 | -0.37 (-0.45/-0.29) | 0.78 | 0.00 | 0.00 | -1.15 | -0.37 |
| 2019 | Popcorn | 0.95 | -1.06 | -0.11 (-0.20/-0.02) | 0.95 | 0.76 | 0.91 | -1.06 | 1.56 |
| Mean |  | 0.87 | -1.11 | -0.24 (-0.33/-0.16) | 0.87 | 0.38 | 0.46 | -1.11 | 0.59 |
| 2018-2019 | Wheat | 1.25 | -1.38 | -0.13 (-0.25/-0.01) | 1.25 | 0.00 | 0.00 | -1.47 | -0.22 |
| 2020 | Popcorn | 0.92 | -1.28 | -0.36 (-0.43/-0.28) | 0.92 | 1.71 | 1.39 | -1.35 | 2.67 |
| Mean |  | 1.09 | -1.33 | -0.24 (-0.34/-0.15) | 1.09 | 0.86 | 0.69 | -1.41 | 1.23 |

### 3.2 Climate change impact of crop rotations

The conservation crop rotation had higher GHG emissions than the conventional crop rotation (15% higher in 2017-2019; 25% in 2018-2020) and also sequestered much more C (−347% as much in 2017-2019; -608% in 2018-2020) (Fig. 1). Consequently, conventional rotations had positive climate change impacts (7440 kg CO_2_-eq./ha in 2017-2019 and 8027 kg CO_2_-eq./ha in 2018-2020), as well as the conservation rotation in 2017-2019 (2422 kg CO_2_-eq./ha). On the other hand, the conservation rotation in 2018-2020 had a negative climate change impact (−688 kg CO_2_-eq./ha), because C sequestration was superior to GHG emissions. For the conventional crop rotations, direct emissions contributed most to the climate change impact. For the conservation crop rotation, direct emissions had a similar magnitude as those of conventional crop rotations, but C sequestered contributed most to the climate change impact.

**Figure 1.**
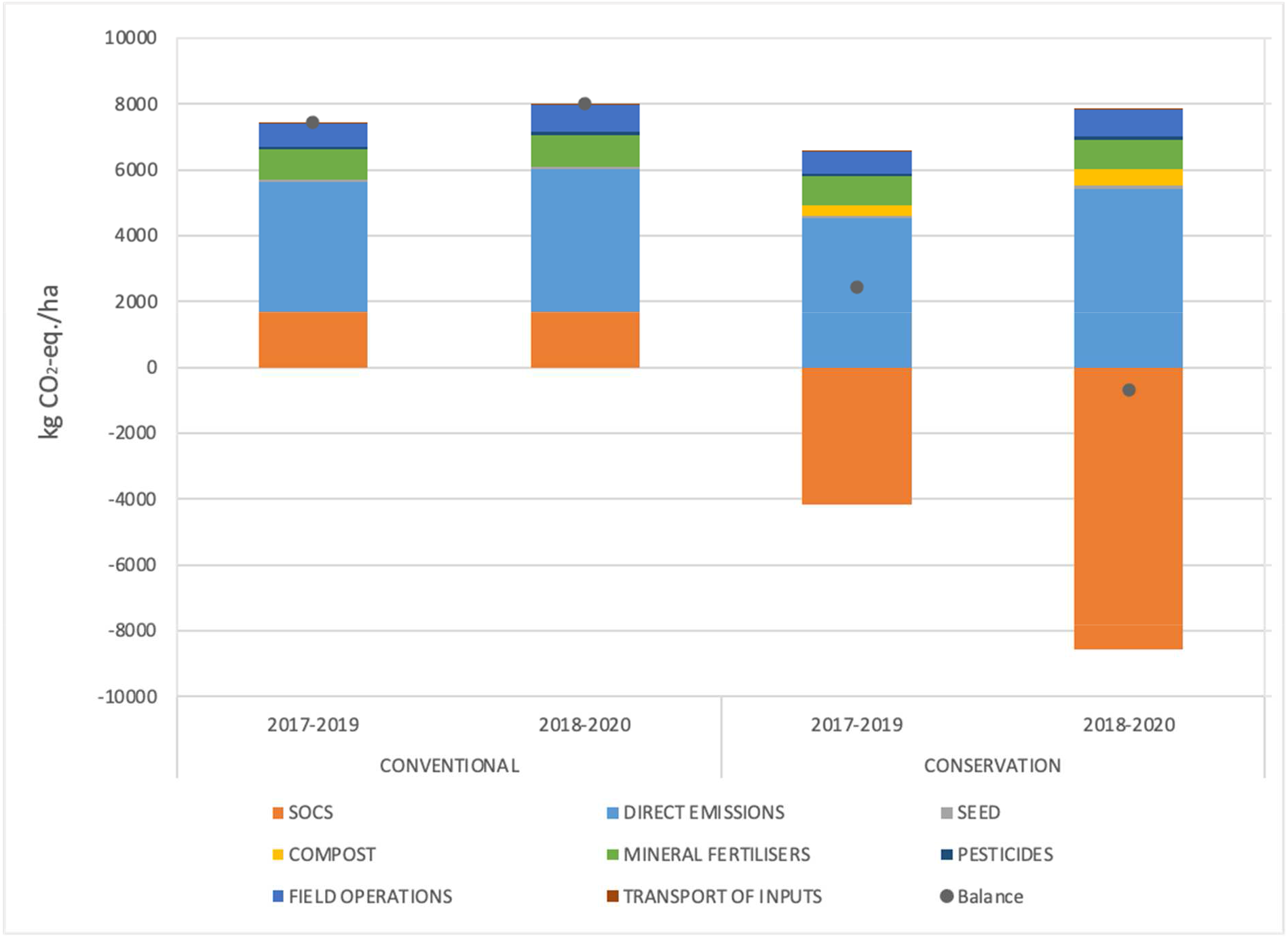
Contribution of soil organic carbon, direct emissions, seed, compost, mineral fertilisers, pesticides, field operations (including irrigation), and transport of inputs to the climate change impact for the two-year popcorn and wheat crop rotation in 2017-2019 and 2018-2020, for conventional and conservation rotations. Dots indicate net impact, which is equal to C sequestration minus greenhouse gas emissions. The inputs and direct emissions of the cover crops were attributed to the popcorn crops.

When averaged over the two crop rotations, the climate change impact considering only emissions was 1% lower for wheat and 8% higher for popcorn for conservation agriculture than for conventional agriculture (Fig. 2). Expressed in kg CO_2_-eq./ha, impacts were 2629 (conventional) and 2598 (conservation) for wheat, 3407 (conventional) and 3680 (conservation) for popcorn, and 966 for cover crops.

**Figure 2.**
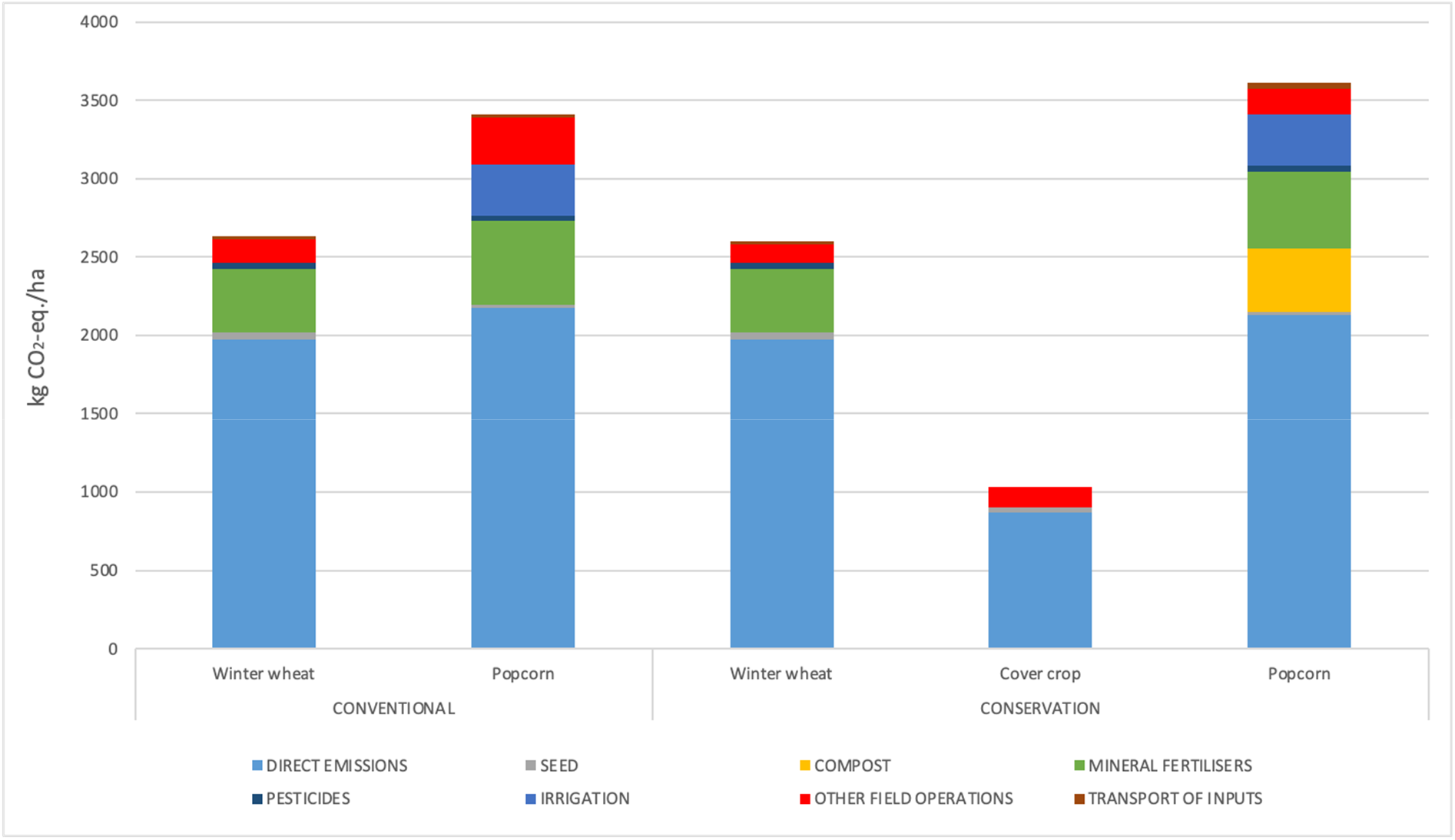
Contribution of direct emissions, seed, compost, mineral fertilisers, pesticides, irrigation, other field operations, and transport of inputs to the climate change impact for individual crops. Values are means for 2017-2018 and 2018-2019 for wheat, for 2019 and 2020 for popcorn, and for 2018-2019 and 2019-2020 for cover crops.

For conservation popcorn, climate change impacts for other field operations and mineral fertilisers were 45% and 10% lower, respectively, than those for conventional popcorn. Transport of inputs and pesticides, which contributed little (ca. 1%) to impacts for all crops, were 78% and 16% higher respectively, for conservation popcorn than for conventional popcorn. The processes that contributed most were direct emissions (64% conventional and 59% conservation), mineral fertiliser (16% conventional and 13% conservation), irrigation (10% conventional and 9% conservation), other field operations (9% conventional and 4% conservation), and compost 11% (only for conservation).

For conservation wheat, climate change impacts for direct emissions and other field operations were similar to and 22% lower, respectively, than those for conventional wheat. Impacts associated with mineral fertilisers, pesticides, and transport of inputs did not differ for conservation and conventional wheat because they used identical amounts. The processes that contributed most were direct emissions (75% conventional and 76% conservation), mineral fertiliser (15% for both conventional and conservation), and other field operations (6% conventional and 4% conservation). For the cover crops, the processes that contributed most were direct emissions (84%), other field operations (13%), and seeds (3%).

### 3.3 Multi-impact comparison of crop rotations

Impacts of the conventional and conservation rotations differed (Table 4). The conservation rotation had lower impacts than the conventional rotation for climate change (−89%) and marine eutrophication (−7%). A 10% decrease or increase of assumed yields of the crops in the conventional rotation resulted in a moderate (−97%/-82%) variation of the difference for the climate change impact (Table 4). The conservation rotation had somewhat higher impacts than the conventional rotation for terrestrial acidification (+9%), land competition (+3%), and cumulative energy demand (+2%).

**Table 4.**
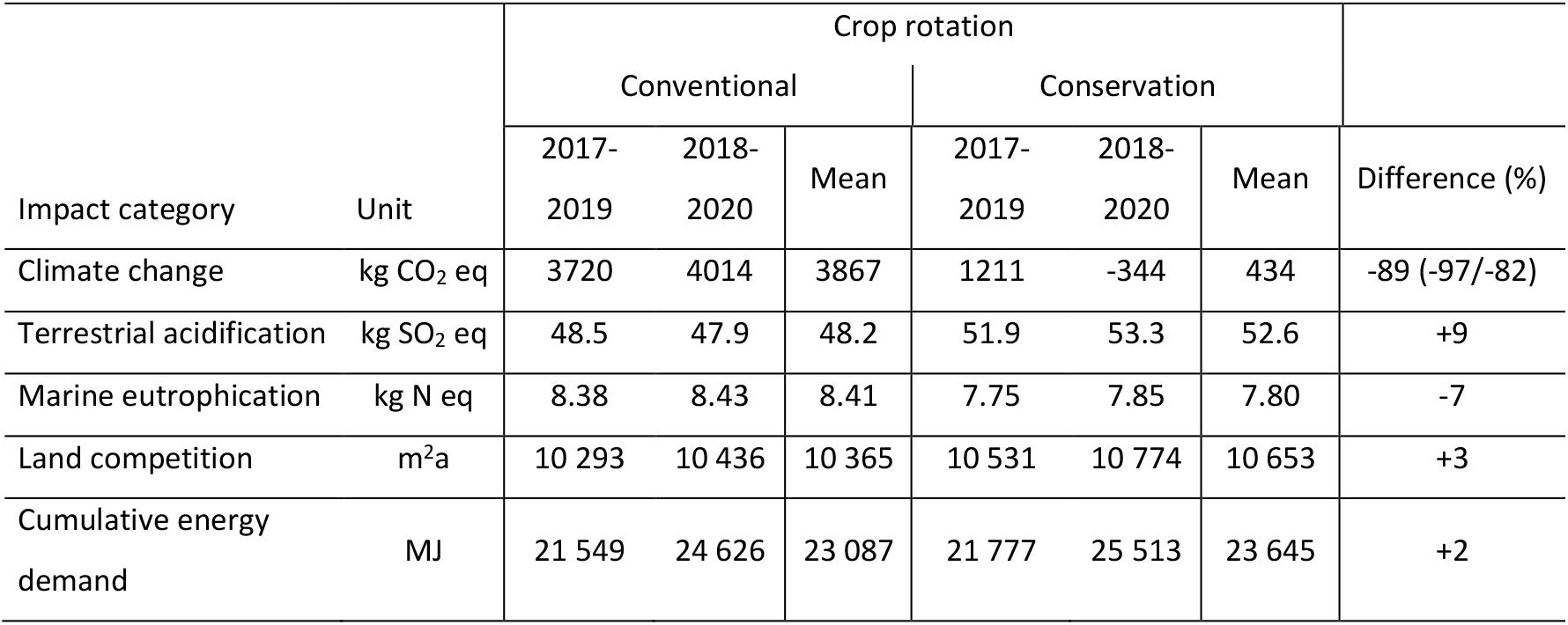
Impacts of conventional and conservation crop rotations (2017-2019 and 2019-2020) for the land management (ha.year) functional unit. Difference expresses impacts of the conservation rotation relative to those of the conventional rotation. Values in brackets for Difference correspond to the effects of a 10% increase or decrease of the conventional crop yields.

When comparing the 2017-2019 and 2018-2020 crop rotations, differences were small, except for climate change for the conservation rotation, which had much lower impacts for 2018-2020 than for 2017-2019 (−169%). Cumulative energy demand was 17% higher for 2018-2020 than for 2017-2019 for both the conservation and conventional systems.

### 3.4 Change in soil organic C stock over time of scenarios

According to the SIMEOS-AMG simulation tool, initial SOC stock was 38.4 t/ha for the three scenarios (Fig. 3). Over time, SOC stock decreased in the Conv scenario but increased in the Cons_CC and Cons_full scenarios. After 100 years, SOC had not completely reached equilibrium. When 95% of the change in SOC stock at 100 years was reached, the subsequent annual change was near zero for each scenario: -0.0005% (Conv), 0.0005% (Cons_CC), and 0.0009% (Cons_full). The Conv scenario reached this 95% threshold after 31 years, losing 9% of the initial SOC stock, whereas Cons_CC and Cons_full reached it after 34 years, gaining 14% and 26% of the initial SOC stock, respectively. After 100 years, estimated SOC stocks were 35.7 (Conv), 43.9 (Cons_CC), and 48.8 (Cons_full) t C/ha.

**Figure 3.**
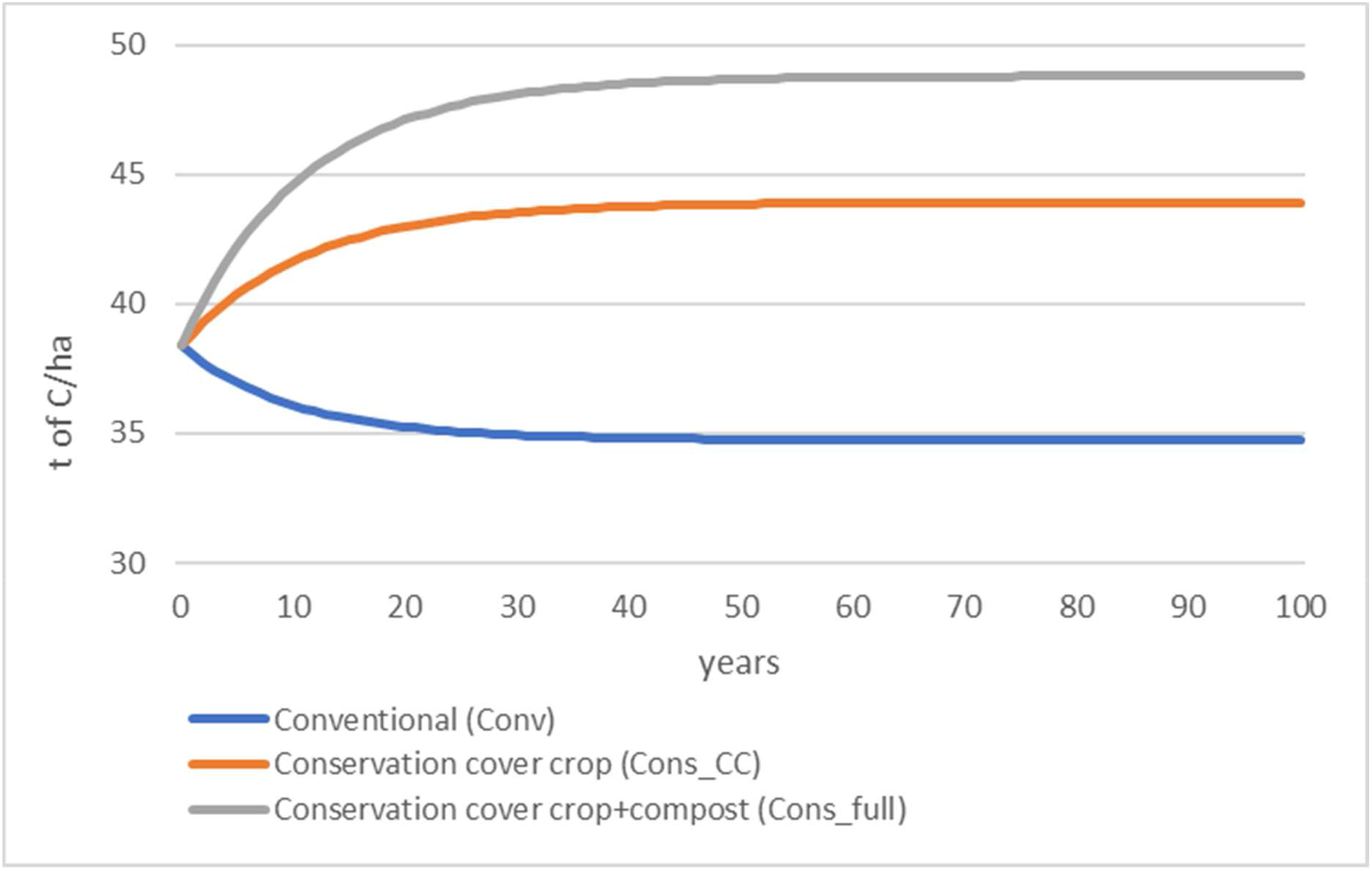
Soil organic carbon stock (0-30 cm) over 100 years according to three scenarios of popcorn and wheat crop rotations as estimated by the SIMEOS-AMG simulation tool. See Table 1 for a description of the scenarios.

In the south west of France approximately 7500 ha of popcorn are grown each year. To quantify its large-scale C sequestration potential we assume that all of this popcorn is grown in a popcorn-wheat rotation which would transition from conventional to conservation agriculture according to the Cons_CC scenario. Over a 34-year period this would allow the sequestration of 43.9 – 38.4 = 5.5 t/ha on 15000 ha (twice the popcorn area, because popcorn is present one year out of two in the rotation. This corresponds to an average annual C sequestration of (5.5 * 15000)/34 = 2426 t. The annual C footprint of a French citizen being 9 t CO_2_-eq. (SDES, 2022), this corresponds to the C footprint of 270 citizens.

Initial annual climate change impacts for the scenarios were 0.19 (Cons_full), 1.64 (Cons_CC), and 4.31 (Conv) t CO_2_-eq./ha. As mentioned, the Cons_full scenario included a sorghum cover crop followed by a fava bean + phacelia cover crop and 6.5 t/ha of compost (Table 1). The Cons_full scenario resembled the 2017-2019 conservation system (Table 3), but differed in that it had a single cover crop (fava bean

+ phacelia) and slightly different crop yields (6.8 and 6.3 t/ha for wheat and popcorn, respectively vs. 5.9 and 6.5 t/ha, respectively in the 2017-2019 system). Assuming constant management practices, the climate change impact decreased over time for Conv but increased for Cons_CC and Cons_full. Climate change impacts for the three systems intersected after 23-28 years, when they became larger for Cons_CC and Cons_full than for Conv.

## 4. Discussion

### 4.1 C inputs and sequestration

According to field data for the Nataïs conservation agriculture system, C input from cover crops was

0.8 and 1.7 t/ha in 2019 (fava bean) and 2020 (sorghum followed by fava bean), respectively (Table 3). The 2020 value is close to the mean C input by cover crops (1.9 t/ha) estimated from the meta-analysis of Poeplau and Don (2015). Considering both C inputs by crop residues and cover crops as well as mineralisation of soil C, mean annual C sequestration for the cropping system was 0.6 and 1.2 t/ha for 2017-2019 and 2018-2020, respectively.

According to the meta-analysis of Sun et al. (2020), the difference in mean annual SOC sequestration of conservation vs. conventional agriculture was 0.35 t C/ha. Conservation agriculture practices do not include organic fertilisation, so for comparison with our results, the contribution of compost to SOC sequestration (0.46 (2019) and 0.69 (2020) t C/ha) should be subtracted. This yields differences in annual C sequestration of conservation vs. conventional system of 0.37 (i.e. 0.83 − 0.46) and 0.78 (i.e. 1.47 − 0.69) t/ha for 2019 and 2020, respectively. When a single cover crop was present, our results thus agreed with those from the meta-analysis, and when two consecutive cover crops were grown, they were twice as high.

In the Nataïs conservation agriculture system, compost contributed 34% (2019) and 35% (2020) to the flow of C to the soil, and was thus a major contributor to the system’s climate change impact (Table 3). Based on composting cost and compost price, we allocated 7.2% of impacts of the composting process to the compost. As composting cost and compost price can vary, a sensitivity analysis was conducted. A 50% decrease or 50% increase in the allocation percentage had a small effect (−5% and + 5%, respectively) on the climate change impact of conservation popcorn (Fig. 5). However, when 100% of the impact of the composting process was allocated to the compost, the climate change impact increased by 140%.

The French low-C label (Soenen et al., 2021) developed a method to quantify SOC sequestration and reductions in GHG emissions. This method uses the AGRIBALYSE LCI of compost of green waste, which allocates 100% of the impact of the composting process to the compost. The climate change impact of this compost is 694 kg CO_2_-eq./t. In our study, compost of green waste contained 172 kg C/t, 41% of which (i.e. 70.5 kg, corresponding to 259 kg CO_2_-eq.) was estimated to be sequestered (Pellerin et al. (2019). Consequently, using this method, applying 1 t of compost of green waste increases the climate change impact of a given crop by 435 kg CO_2_-eq. (i.e. 694 – 259), not counting emissions associated with applying compost. This example illustrates the importance of allocation choices on potential climate change impacts, and the need to establish a meaningful consensus on this.

### 4.2 SOC dynamics and climate change impact

#### 4.2.1 SOC dynamics

In their recent review of 106 mainly short-term (2-3 year) experiments on the impact of cover crops, Abdalla et al. (2019) found mean annual C sequestration of 0.54 t/ha. For our Cons_CC scenario, the short-term (2-year) annual C sequestration was 0.57 t/ha, close to their value.

#### 4.2.2 Climate change impact

Over time, the soil was a C sink in the conservation scenarios but a C source in the conventional scenario (Fig. 3). Considering both GHG emissions and changes in SOC stock, the net climate change impact of the conservation scenarios increased over time, eventually overtaking that of the conventional scenario (Fig. 4). This was due to the slightly higher GHG emissions of the conservation scenarios and the gradual decrease in the amount of SOC sequestered by the conservation scenarios and released by the conventional scenario. This illustrates that conservation agriculture can sequester a large amount of C until the SOC stock reaches a new equilibrium is after a few decades.

**Figure 4.**
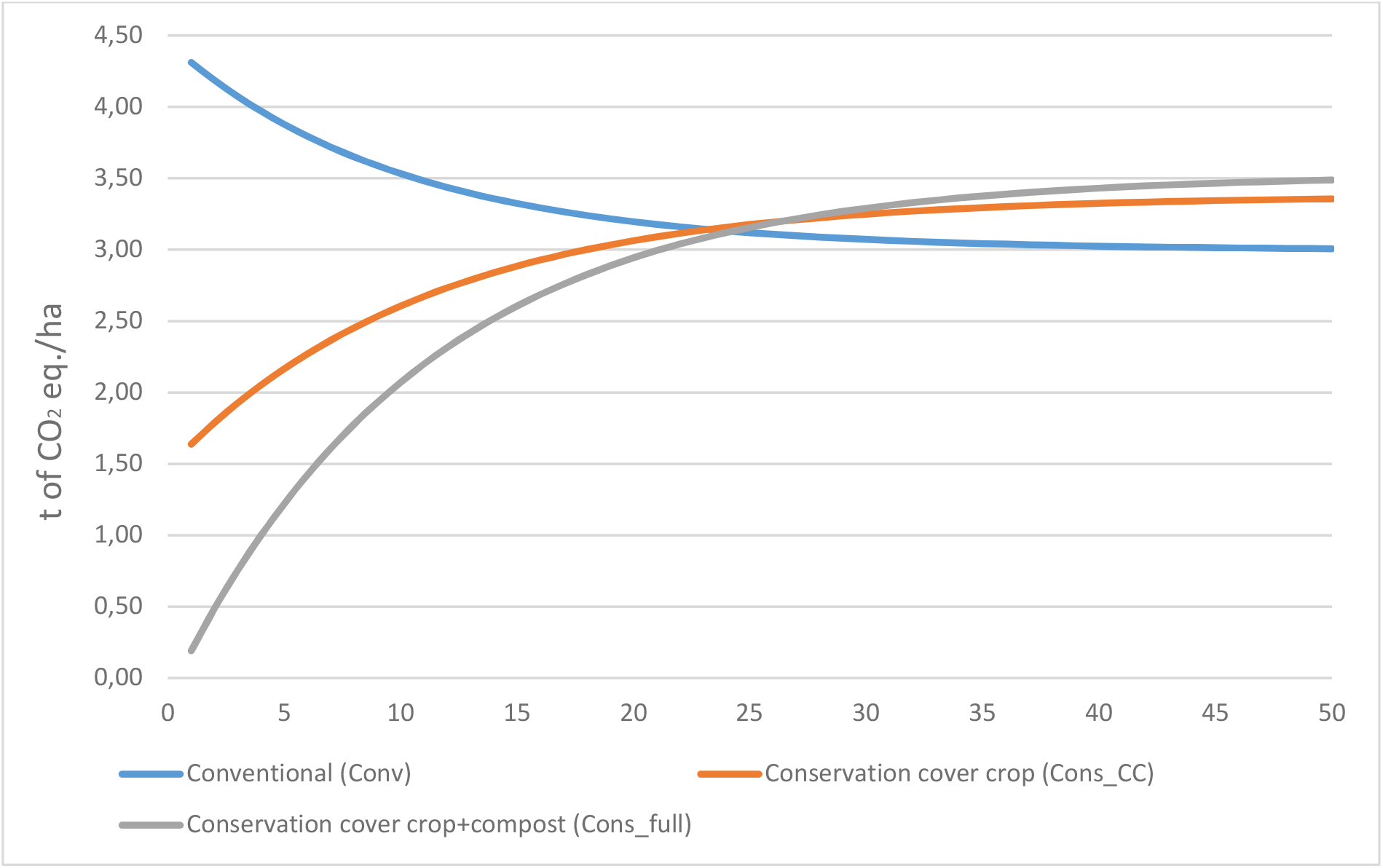
Net climate change impact over 50 years considering greenhouse gas emissions and soil organic carbon sequestration according to three scenarios of popcorn and wheat crop rotations. See Table 1 for a description of the scenarios.

**Figure 5.**
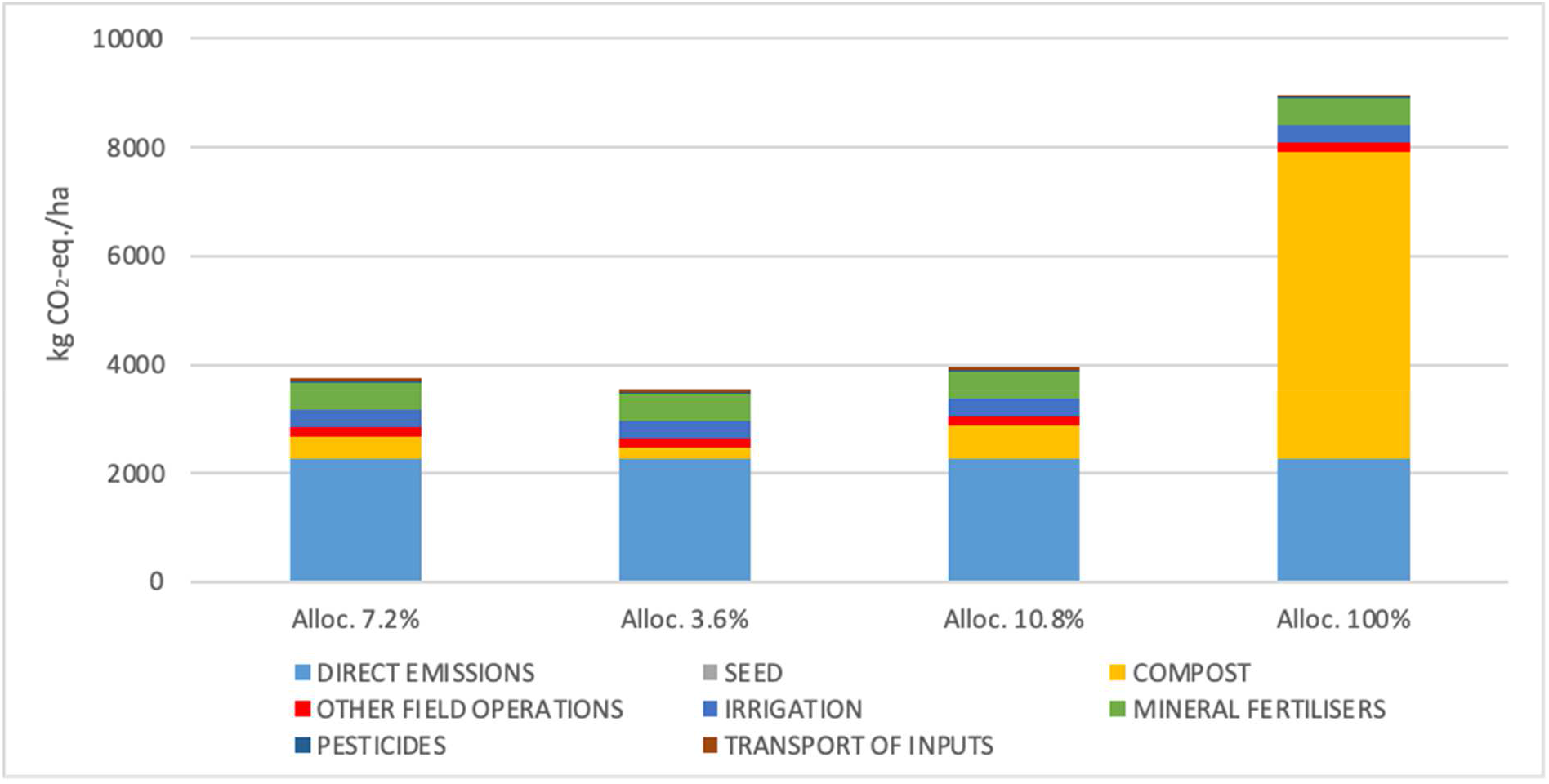
Contribution of direct emissions, seed, compost, other field operations, irrigation, mineral fertilisers, pesticides and transport of inputs to the climate change impact of conservation agriculture popcorn as a function of the percentage of impact allocated to compost: 7.2% (this study), 3.6% (50% decrease), 10.8% (50% increase), and 100% (original life cycle inventory from AGRIBALYSE; Avadi (2020)).

It would be possible to increase C sequestration of the conservation scenario further by increasing C inputs, for instance by shifting towards more complex systems such as agroforestry (Lorenz and Lal, 2014). In their review, Cardinael et al. (2018) indicate that converting an arable system to a silvo-arable system yielded mean annual C sequestration rates of 0.21, 0.28, and 1.12 t/ha in SOC, below-ground biomass, and above-ground biomass, respectively.

### 4.3 Effects of the conservation system on other impact categories

Conventional and conservation agriculture systems differed strongly in their climate change impact, but differences for other impact categories were relatively small. Terrestrial acidification was 9% higher for conservation agriculture than for conventional agriculture. This difference was due to NH_3_ emissions, which were higher for the conservation agriculture system, which used compost in addition to mineral fertiliser, whereas the conventional agriculture system used only mineral fertiliser. Marine eutrophication was 7% lower for the conservation system than for the conventional system due to lower NO_3_ leaching because of the cover crop in the former. Land competition was 3% higher for the conservation system than for the conventional system because more land was needed to produce cover crop seeds. Cumulative energy demand was 2% higher for the conservation system than for the conventional system because the cover crops required more inputs (i.e. seeds, diesel, and transport of inputs).

### 4.4 Study limitations

The assessment of short-term effects of conservation agriculture was based on real-life farm data on yield and farming practices for the conservation rotation, while farming practices of the conventional rotation were based on expert judgement and its crop yields were assumed to be identical to those of the conservation rotation. Although the assumption regarding yield level is supported by the literature and we consider the expert judgement on farming practices to be of high quality, the lack of field data on the reference conventional system constitutes a limitation of this study. A 10% decrease or increase of assumed conventional crop yields had a modest effect on C sequestration and climate change impact, suggesting that our results are nevertheless robust.

Furthermore effects of the conservation and conventional agriculture rotations on soil carbon dynamics are based on modelling results. This also constitutes another limitation of this study. However, an evaluation of the AMG model by comparing it to SOC results from 60 long-term field trials in France revealed that the model accurately simulated the changes in SOC stocks over time (Clivot et al, 2019). Thus, in spite of the absence of field data, we consider these model-based results to be sufficiently solid.

### 4.5 Further research

Although SOC sequestration due to cover crops ceases when a new equilibrium is reached, cover crops have other beneficial effects (Blanco-Canqui et al., 2015). Due to their albedo effect, they mitigate climate change by reflecting more sunlight than most bare soils (Kaye and Quemada, 2017; Carrer et al., 2018). The annual albedo effect of cover crops on climate change has been estimated to be equivalent to 120-460 kg CO_2_-eq./ha (Kaye and Quemada, 2017), smaller than the average effect of cover crops on SOC, but not insignificant. Other continued beneficial effects of cover crops include reduction of soil erosion, fixation of atmospheric N, reduction of NO_3_ leaching, weed suppression, and improvement of the structure of soil microbial communities (Blanco-Canqui et al., 2015).

## 5. Conclusions

This study revealed that implementing an innovative conservation agriculture system initially decreased the climate change impact of a popcorn and wheat rotation strongly, while moderately influencing marine eutrophication, terrestrial acidification, land competition, and cumulative energy demand. Scenario modelling revealed that a new SOC equilibrium was nearly reached after ca. 30 years, when C sequestration virtually ceased, and the climate change impact of the conservation scenarios was slightly higher than that of a conventional scenario. Modelling over 100 years revealed that at near soil C equilibrium, the SOC stock of the conventional scenario had decreased by 9%, whereas that of the conservation scenario had increased by 14% and, when compost was used, by 26%.

The climate change impact of the system using compost of green waste depended strongly on how impacts of the composting process were allocated between its waste treatment and compost production functions. This finding illustrates the need to establish a consensus on the allocation of composting impacts.

Conservation agriculture can sequester a large amount of C until the SOC stock reaches a new equilibrium after a few decades. Shifting towards more complex systems such as agroforestry could enhance C sequestration further. Cover crops have other continuing beneficial effects such as climate change mitigation by reflecting more sunlight than bare soil, reduction of soil erosion, fixation of atmospheric N, reduction of NO_3_ leaching, and weed suppression.

## Acronyms and notations

C: carbon
Ccomp: organic C input from compost
Ccover: organic C input from cover crops
Ccrop: organic C input from cash-crop residues
Cd: cadmium
Cmin: mineralisation of soil C
Cr: chrome
Cseq: organic C input from humified C sequestered
Cu: copper
°C: degree Celsius
CO_2_: carbon dioxide
CO_2_-eq.: carbon dioxide-equivalent
Cons_CC: conservation agriculture with cover crops scenario
Cons_full: conservation agriculture with cover crops and compost scenario
Conv: conventional agriculture scenario
GHG: greenhouse gas
Ha: hectare
Hg: mercury
IPCC: Intergovernmental Panel on Climate Change
ISO: international organization for standardization
LCA: life cycle assessment
LCI: life cycle inventory
N: nitrogen
NH_3_: ammonia
Ni: nickel
NO_3_: nitrate
NO_x_: nitrogen oxides
N_2_O: nitrous oxide
P: phosphorous
Pb: lead
PO_4_: phosphate
SOC: soil organic carbon
t: ton
Zn: zinc

## CRediT authorship contribution statement

Maria Vittoria Guidoboni: Methodology, Formal analysis, Writing – original draft. Annie Duparque: Methodology, Formal analysis, Writing-Reviewing and Editing. Joachim Boissy: Formal analysis, Methodology. Jean-Christophe Mouny: software, formal analysis. Julie Auberger: Software, Data curation, Methodology, Writing-Reviewing and Editing. Hayo van der Werf: Conceptualisation, Methodology, Writing-Reviewing and Editing, Supervision.

## Acknowledgements

We thank the Nataïs team for its support, in particular Anne-Marie Joliet, who answered our many questions during data collection. We thank Eric Ceschia of CESBIO/INRAE for his reactivity and help, as well as Sylvain Hypolite (Agro d’Oc) for his expert knowledge. We thank our colleagues of the Carbon Farming project, especially Gécica Yogo for her coordination role. This project received funding from the Carbon Farming research and innovation project under grant agreement no. TCL 210065 and was supported by EIT Climate-KIC.

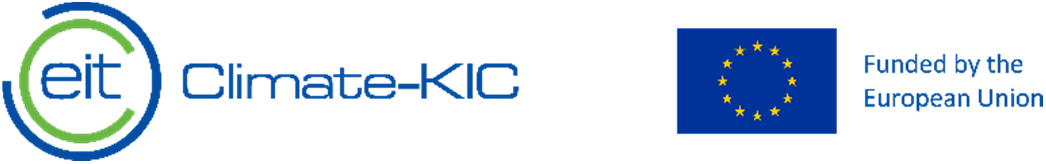

---

**Table S1:**
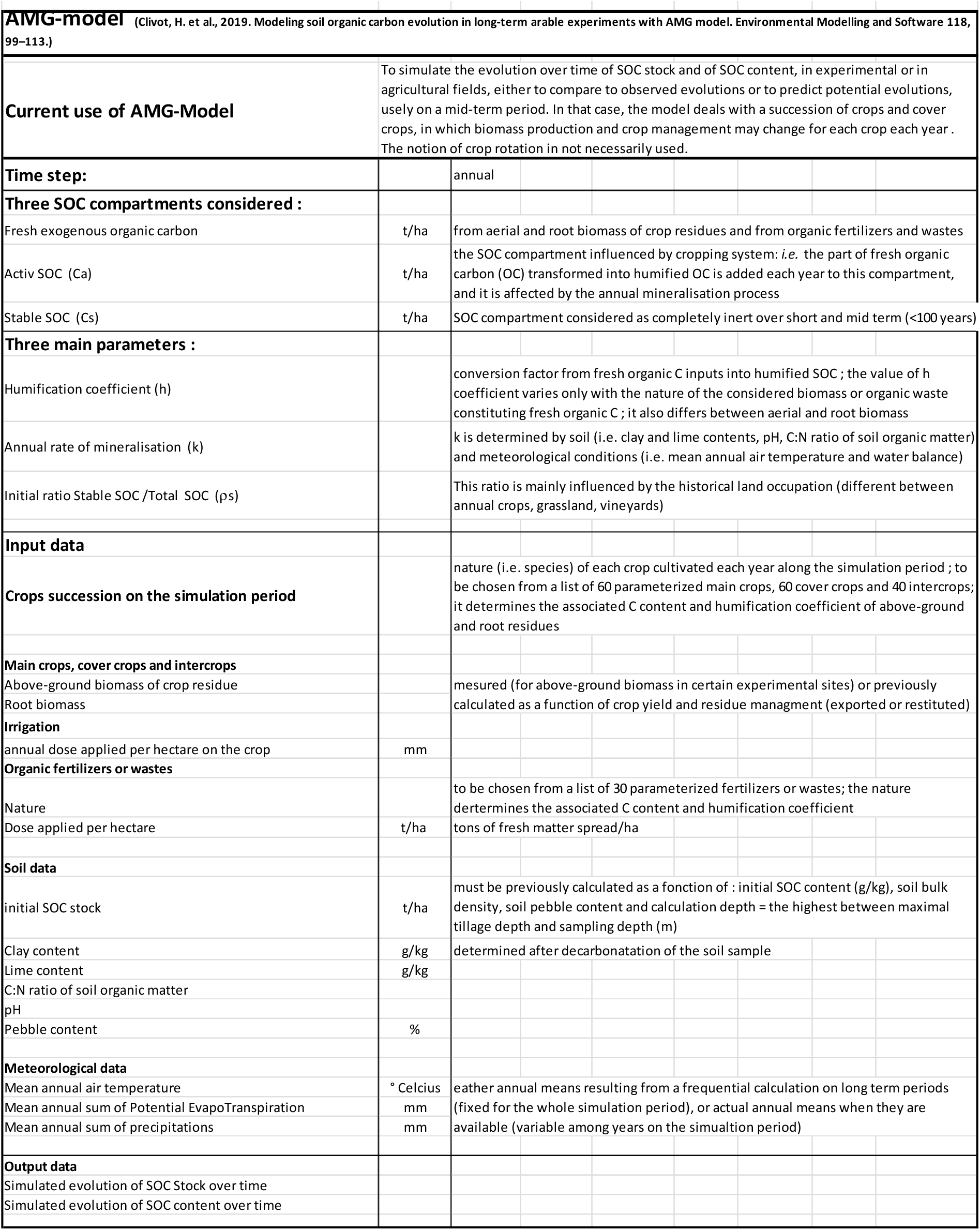
Description of AMG model (cf. calculation of short-term soil organic carbon (SOC) stock variations based on field data and expert judge

**Table S2:**
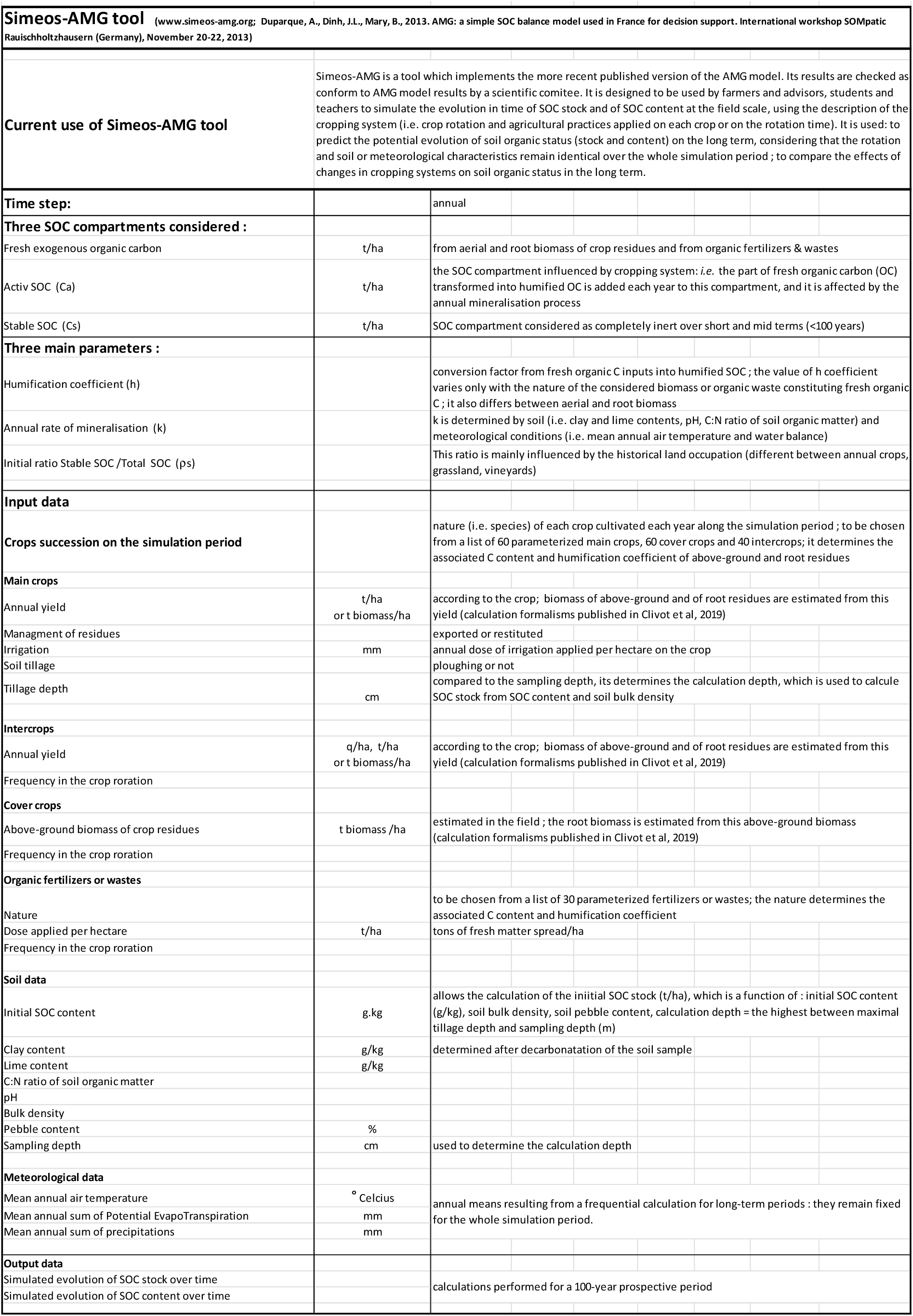
Description of Simeos-AMG tool (cf. calculation of long-term soil organic carbon (SOC) stock variations for the three prospective scenarios)

